# Comprehensive Multi-omics Analysis Reveals the Core Role of Glycerophospholipid Metabolism in Rheumatoid Arthritis Development

**DOI:** 10.1101/2023.02.15.528612

**Authors:** Congcong Jian, Lingli Wei, Tong Wu, Shilin Li, Tingting Wang, Jianghua Chen, Chengjia Chang, Jie Zhang, Binhan He, Jianhong Wu, Jiang Su, Jing Zhu, Min Wu, Yan Zhang, Fanxin Zeng

**Author notes:** These authors contributed equally to this paper (Congcong Jian, Lingli Wei, Tong Wu). Corresponding Author: Min Wu, Ph.D. Huaxi MR Research Center (HMRRC), Department of Radiology, West China Hospital of Sichuan University, Chengdu, 610041, China., Yan Zhang, M.D, Ph.D. Lung Cancer Center of West China Hospital, Sichuan University, Chengdu, Sichuan, China., Fanxin Zeng, Ph.D. School of Basic Medical Science, Chengdu University of Traditional Chinese Medicine, Chengdu, China; Department of Clinical Research Center, Dazhou Central Hospital, Dazhou, Sichuan, China; Department of Big Data and Biomedical AI, College of Future Technology, Peking University Beijing 100871, China.

## Abstract

**Objectives:** Rheumatoid arthritis (RA) is a chronic autoimmune disease with complex causes and recurrent attacks that can easily develop into chronic arthritis. Our study aims to elucidate potential mechanism among control, new-onset RA (NORA) and chronic RA (CRA) with multi-omics analysis.

**Methods:** A total of 162 subjects were included in our study, 121 subjects were used for 16S rRNA, ITS sequencing and metabolomics analysis and 41 subjects were used for transcriptomics analysis. Enrichment analysis was based on significant difference metabolites and genes. Protein-protein interaction network and correlation analyses were performed to interpret the interactions among intestinal flora, metabolites and genes. We applied three models to distinguish between NORA and CRA using computational OR values, LASSO and random forest, respectively.

**Results:** Our results demonstrated intestinal flora disturbance in RA development, with significantly increased abundance of *Escherichia-Shigella* and *Proteobacteria* in NORA. We also found that the diversity was significantly reduced in CRA compared to NORA through fungi analysis. Moreover, we identified 28 differential metabolites between NORA and CRA. Pathway enrichment analysis revealed significant dysregulation of glycerophospholipid metabolism and phenylalanine metabolism pathways in RA patients. Next, we identified 40 differentially expressed genes between NORA and CRA, which acetylcholinesterase (ACHE) was the core gene and significantly enriched in glycerophospholipid metabolism pathway. Correlation analysis showed a strong negatively correlation between glycerophosphocholine and inflammatory characteristics.

**Conclusions:** These findings revealed that glycerophospholipid metabolism plays a crucial role in the development and progression of RA, providing new ideas for early clinical diagnosis and optimizing treatment strategies.

## Introduction

Rheumatoid arthritis (RA) is a chronic, systemic autoimmune disease, characterized by symmetrical synovial inflammation and eventual involvement of other organ systems (1–3). According to epidemiological surveys, the global prevalence of RA is 0.2%-1.0%, and nearly 5 million people in China suffer from RA, with a prevalence of 0.28%-0.41% (4). The pathogenesis of RA is not fully understood, and its highly specific, inherited and environmental factors combine to influence the onset and progression of the disease (5–7). Symptoms of new-onset RA (NORA) are mainly characterized by high disease activity, joint inflammation and pain. Most patients have delayed treatment due to ineffective treatment regimens or poor compliance, and as the disease progresses and the inflammatory state worsens, the joints and articular cartilage were destroyed, eventually evolving into chronic rheumatoid arthritis (CRA) with a range of extra-articular damages (8, 9). Therefore, early diagnosis and early treatment of RA can effectively prevent disease progression, joint damage and destruction of other organ systems in most patients.

Among the many influential factors, intestinal flora is considered to be an important trigger for immune system abnormalities in RA (10). Previous studies showed that a decrease in the composition and diversity of the intestinal flora in RA patients, with an increase in the abundance of *Klebsiella, Escherichia* and a decrease in the abundance of *Megamonas* and *Enterococcus,* and an expansion of *Prevotella* associated with an increased susceptibility to arthritis, suggesting that the development of RA is closely associated with dysbiosis of the intestinal flora (11, 12). In recent years, with the development of metabolomics technology, more and more evidences revealed that patients with RA have significant changes in plasma metabolites and metabolic pathways, such as lipid metabolism and amino acid metabolism (11, 13). Transcriptomic analysis is widely used in the field of RA research. Gene expression profiles of peripheral blood mononuclear cells (PBMCs) could reveal the pathological process and pathogenesis of RA involving immune cells, and play an important role in predicting response to drug therapy, screening for key differential genes and explaining the pathogenesis of RA (14, 15). A study had demonstrated that HLA-DRB1 was a susceptibility gene that triggers RA, affecting disease activity and treatment response (16).At present, multi-omics combined analysis is broadly applied in the research field, which can explore the interactions and potential links between gut microbes and metabolites, and elucidate the pathogenesis of diseases as a result of the combined influence of multiple factors. However, multi-omics researches are less well studied in explaining the pathogenesis of RA development, and it is essential to use multi-omics approaches to gain a comprehensive understanding of the pathogenesis of RA.

In our study, we combine gut flora, plasma metabolism and transcriptome analysis to explore the changes and potential relationships between control, NORA and CRA patients, to elucidate the interactions between gut microbes, plasma metabolites and genes, and to reveal the pathogenesis of RA with the joint influence of multiple factors, providing a new perspective and research direction for early clinical diagnosis and precise treatment.

## Materials and methods

### Participant recruitment

The study population consisted of RA patients aged >18 years from the Department of Rheumatology and Immunology, and healthy control from the medical examination center, Dazhou Central Hospital. NORA were included those with less than 6 months of disease and who have never used antirheumatic drugs, CRA with more than 6 months of disease who have used traditional antirheumatic drugs or biologics. Patients were diagnosed with rheumatoid arthritis in this study according to the 2010 American College of Rheumatology (ACR) and European League of Rheumatology (EULAR) criteria (17). Clinical data from RA patients were recorded, including rheumatoid factor (RF), erythrocyte sedimentation rate (ESR), C-reactive protein (CRP), joint tenderness number of 28 joints (TJC28), joint swelling number of 28 joints (SJC28), disease activity score of 28 joints (DAS28), and interleukin-6 (IL-6). This study was approved by the Medical Ethics Review Board of Dazhou Center Hospital (ID Number:022/2021), and all the subjects signed informed consent.

### Study design

Our study included 162 subjects (NORA=34, CRA=53, control=75), fecal and blood samples were collected from 121 subjects for 16S, ITS sequencing and metabolomics analysis respectively, to identify differential flora and differential metabolites between control, NORA and CRA. PBMCs were collected from 41 subjects for transcriptome sequencing to identify differentially expressed genes among three groups, and finally, a combined multi-omics analysis and receiver operating characteristic (ROC) analysis was performed to establish classification models, details as described in Figure 1.

**Figure 1.**
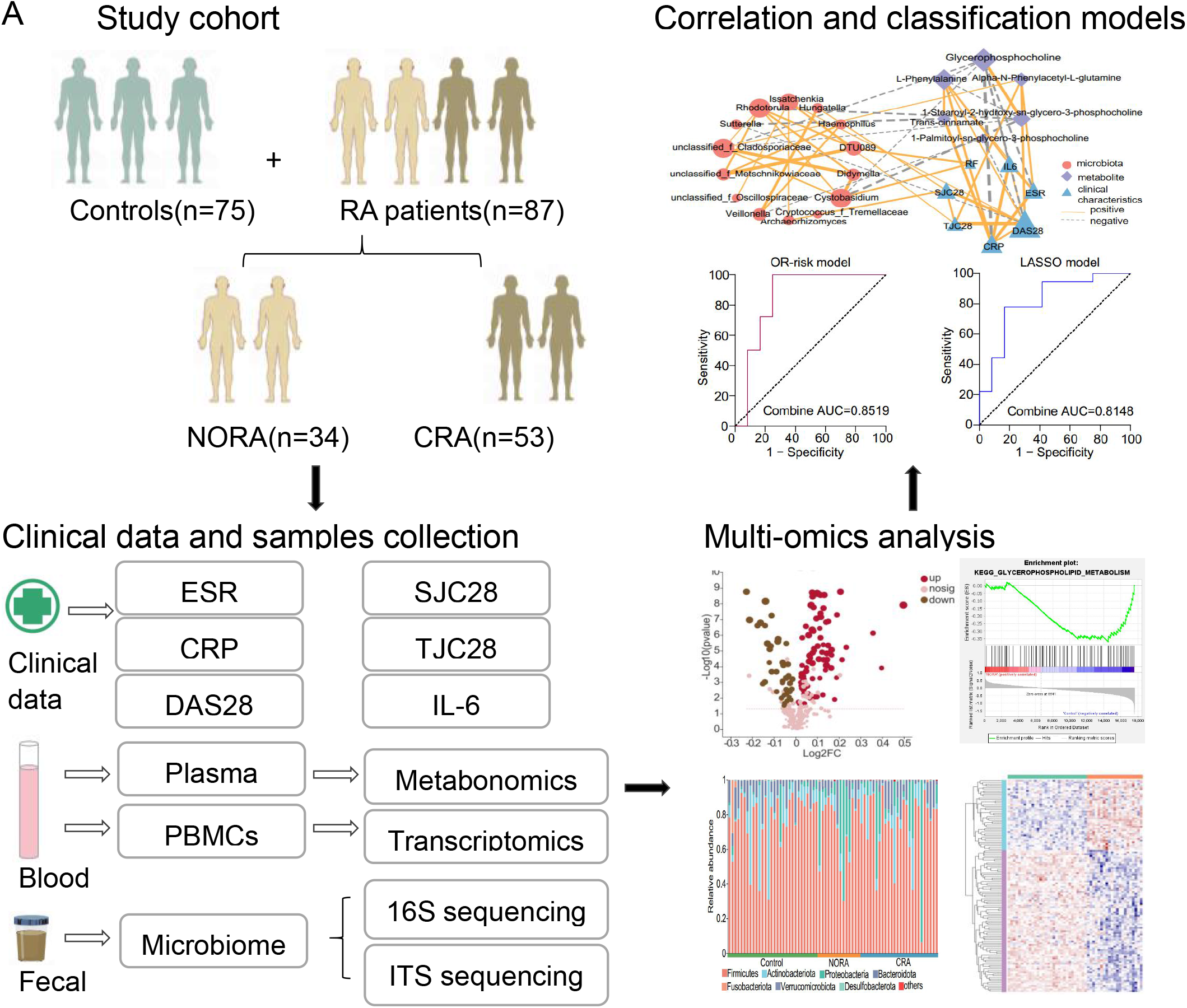
Overview of the study design. NORA, new-onset rheumatoid arthritis; CRA, chronic rheumatoid arthritis; ESR, erythrocyte sedimentation rate; CRP, C-reactive protein; DAS28, disease activity score; SJC28, number of swollen joints; TJC28, number of tender joints; IL-6, interleukin-6; PBMCs, peripheral blood mononuclear cells.

### Intestinal microbiomics analysis

16S rRNA and ITS sequencing on stool samples from subjects were performed. We sampled the amplicon sequence variants (ASVs) by minimum number of sequences, and subsequently we performed gut microbial community composition analysis, alpha diversity and beta diversity analysis for each group of samples. Details as described in supplementary methods.

### Metabolomic analysis

We collected plasma samples from subjects for non-targeted metabolomics assays, and the raw data were quality controlled, corrected and logarithmically processed to obtain data matrix, and we performed orthogonal partial least squares discriminant analysis (OPLS-DA) to identify differential metabolites (VIP>1, p<0.05) and to identify differential metabolite enriched pathways.

### Transcriptomic analysis

We performed transcriptomic sequencing of PBMCs samples, and the data were rigorously processed for selection and identification of differential genes, followed by Kyoto Encyclopedia of Genes and Genomes (KEGG) and Gene Ontology (GO) pathway enrichment analysis.

### Multi-omics combination and ROC analysis

We used protein-protein interaction analysis and correlation network to reveal potential relationships between gut microbes, plasma metabolites and genes. Finally, we used three algorithms of OR value, LASSO and random forest to select crucial features to build classification models for NORA and CRA, and ROC curves were used to evaluate the performance of the classification models.

## Results

### Study population

In our study, we found that SJC28 and the level of IL-6 were higher in NORA than in CRA patients and were apparently significant. We also noticed that CRP, TJC28 and DAS28 were higher in NORA compared to CRA, although there was no difference between them. Similarly, the levels of ESR and RF were not different between two groups, as detailed in Supplementary Table 1.

### Significant dysregulation of metabolites and metabolism pathways in the plasma metabolic profile of RA patients

To identify differential metabolites between control, NORA and CRA groups, we performed non-target metabolomics analysis of three groups. we found 124 differential metabolites between NORA and control, of which 72 were anionic and 52 were cationic metabolites. Orthogonal partial least squares discriminant analysis (OPLS-DA) demonstrated a clear separation of differential metabolites between two groups in the anionic and cationic modes (Supplementary Figure 1A-B). A volcano plot showed that 124 differentially significant metabolites, with 83 up-regulated and 41 down-regulated metabolites significant metabolites (Supplementary Figure 1C). Meanwhile, we identified 114 differential metabolites between CRA and control, and OPLS-DA plots illustrated a clear distinction between differential metabolites in anionic and cationic modes, respectively (Supplementary Figure 1D-E), and volcano demonstrated significant upregulation of 40 metabolites and downregulation of 74 metabolites (Supplementary Figure 1F). Furthermore, we also identified 28 differentiated metabolites between NORA and CRA patients, and OPLS-DA plots demonstrated an apparent differentiation between two groups in terms of differential metabolites in anionic and cationic modes (Figure 2A-B). Volcano plot illustrated that 10 differential metabolites were significantly upregulated and 18 differential metabolites were significantly downregulated (Supplementary Figure 1G). To observe the clustering of differential metabolites between groups, the heat maps showed 124, 114, and 28 differential metabolites clearly distinguished between the three groups, respectively (Supplementary Figure 1H-J).

**Figure 2.**
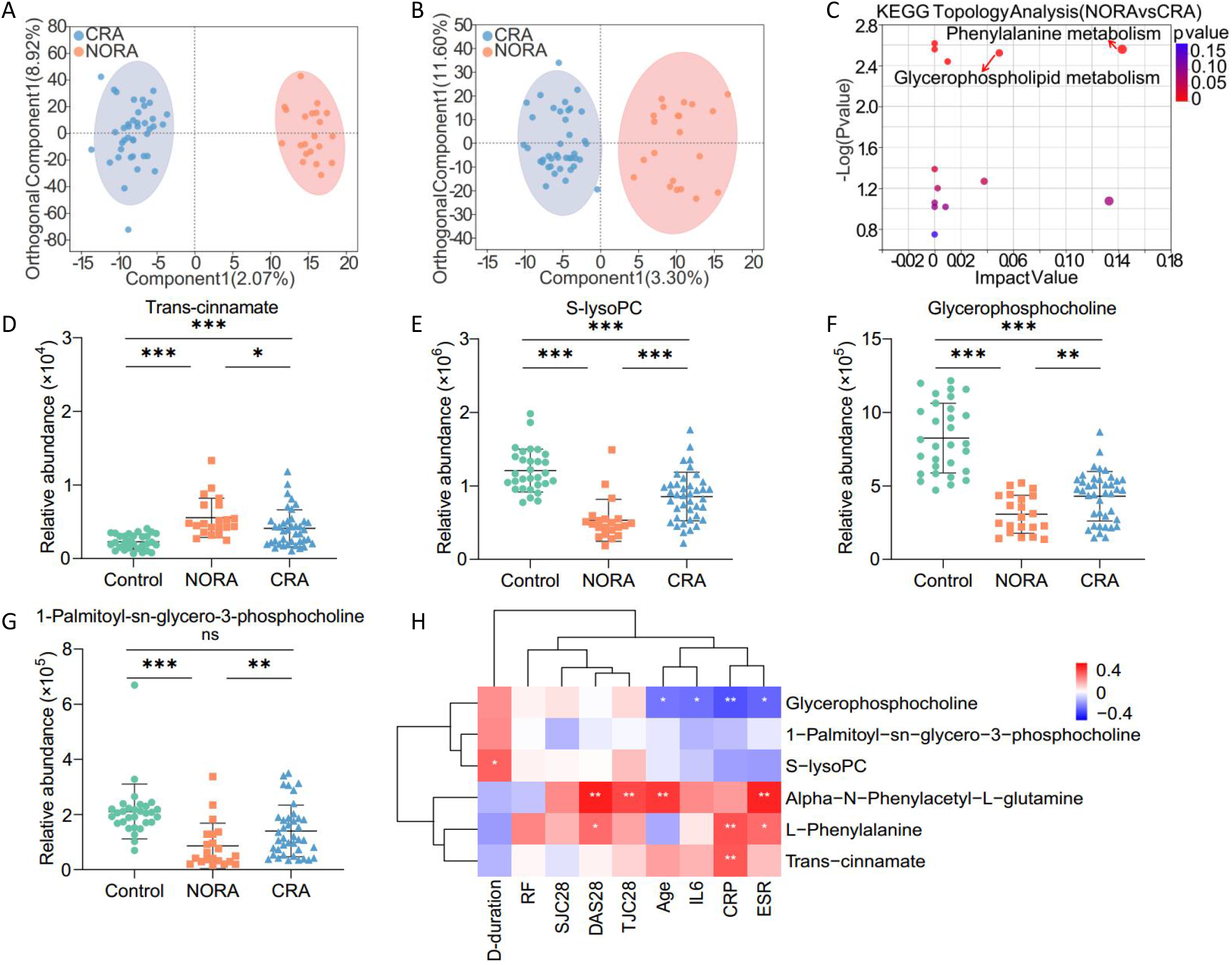
Identification of differential metabolites in plasma metabolic profile between NORA and CRA patients. (A, B) OPLA-DA analysis showed 28 differential metabolites between NORA and CRA in anionic and cationic patterns, respectively. (C) The bubble diagram showed a significant enrichment of differential metabolites between NORA and CRA in the glycerophospholipid metabolism and phenylalanine metabolism pathways, including 1-Palmitoyl-sn-glycero-3-phosphocholine, S-lysoPC, glycerophosphocholine, trans-cinnamate, L-Phenylalanine and alpha-N-Phenylacetyl-L-glutamine. (D-G) Scatter plots demonstrated the relative abundance of 4 differential metabolites on glycerophospholipid metabolism and phenylalanine metabolism pathways between controls, NORA and CRA. (H) Correlation heatmap showed the interrelationship between 6 differential metabolites and clinical characteristics. (S-lysoPC: 1-Stearoyl-2-hydroxy-sn-glycero-3-phosphocholine; D-duration: disease duration).

Afterwards, we performed Kyoto Encyclopedia of Genes and Genomes (KEGG) enrichment analysis of differential metabolites and found 6 metabolic pathways were altered in NORA and 9 metabolic pathways were changed in CRA patients compared to control, with significant deregulation of glycerophospholipid metabolism, tyrosine metabolism, histidine metabolism, citrate cycle (TCA cycle), and glycine, serine, and threonine metabolism pathways common among them (Supplementary Figure 1K-L). We also detected glycerophospholipid metabolism and phenylalanine metabolism pathways were significantly disordered between NORA and CRA patients, including 6 differentially significant metabolites (1-Palmitoyl-sn-glycero-3-phosphocholine, 1-Stearoyl-2-hydroxy-sn-glycero-3-phosphocholine (S-lysoPC), glycerophosphocholine, trans-cinnamate, L-Phenylalanine and alpha-N-Phenylacetyl-L-glutamine) (Figure 2C-G, Supplementary Figure 1M-N, Supplementary Table 2), and 6 differential metabolites were also significantly differentiated between NORA versus control and CRA versus control. More importantly, we found that the expression levels of the first three metabolites were higher in controls than in RA patients, while the last three metabolites were higher in RA patients than in controls. Therefore, we performed correlation analysis of 6 metabolites, and the heatmap demonstrated a significant positively correlation between trans-cinnamate and L-Phenylalanine, and a strong positively association among 1-Palmitoyl-sn-glycero-3-phosphocholine, S-lysoPC, and glycerophosphocholine. While a significant negatively correlation between glycerophosphocholine and trans-cinnamate (Supplementary Figure 1O). Simultaneously, we found strong correlations between six differential metabolites and clinical characteristics, with significant negative correlations between glycerophosphocholine and ESR, CRP, IL-6 and age, positive correlations between S-lysoPC and disease duration. The heat map also exhibited a significant positive correlation between DAS28, CRP and L-Phenylalanine (Figure 2H).

These results suggested significant alterations in plasma metabolic profile of RA patients, with disturbed amino acid metabolism and lipid metabolic pathways, and glycerophospholipid metabolism pathway appeared to play an important role in the progression of RA disease

### An apparently increased abundance of harmful intestinal bacteria and decreased probiotic bacteria in RA patients

To investigate the community structure of intestinal bacteria among control, NORA and CRA patients, we performed 16S rRNA sequencing on stool samples from subjects. We found a reduced number of gut bacteria in NORA compared to control, although there was no significant difference between them, and the Venn diagram showed a total of 133 species between the three groups (Supplementary Figure 2A). In community composition analysis, the relative abundance of the *Proteobacteria* was significantly increased at the phylum level in NORA compared to control, while the abundance of *Firmicutes, Bacteroidota,* and *Actinobacteriota* did not differ between the three groups (Figure 3A-B, Supplementary Figure 2B-D). At the family level, stacked bar chart showed the relative abundance of each family in the three groups in different samples, and we found that *Lachnospiraceae* was the dominant species in three groups (Supplementary Figure 2E). The relative abundance of *Pasteurellaceae* was significantly increased in NORA compared to control and CRA groups (Supplementary Figure 2F). Moreover, we found an apparent increase in the abundance of *Enterobacteriaceae* and decrease in the abundance of *Bifidobacteriaceae* and *Acidaminococcaceae* in NORA patients (Figure 3C-D, Supplementary Figure 2G). We also focused on a reduced abundance of *Bacteroidaceae* in CRA, while *Prevotellaceae* and *Lactobacillaceae* did not differ between the three groups (Supplementary Figure 2H-J). In alpha diversity analysis, we discovered no significant differences in community richness and diversity between control, NORA and CRA patients (Supplementary Figure 3A-F). In beta diversity based on bray-curtis distance, principal co-ordinates analysis (PCoA) showed that NORA could be distinguished from control (P=0.021), whereas no distinction could be observed between CRA and control (P=0.055) and NORA compared with CRA (P=0.535) (Figure 3E, Supplementary Figure 3G-H). Basing on the Wilcoxon rank-sum test, we found 26 differential genera at genus level between controls and NORA, with *Escherichia-Shigella* and *Veillonella* significantly increasing in abundance and *Bifidobacterium* decreasing in NORA (Figure 3F). Meanwhile, we identified 17 differential genera between CRA and controls, with a dramatically increased abundance of *Eubacterium_hallii_group* in CRA, and we also identified 7 differential genera between NORA and CRA, with a significant decrease in abundance of *Veillonella* and *Haemophilus,* and *Anaerostipes* significantly increased in CRA (Figure 3G-H).

**Figure 3.**
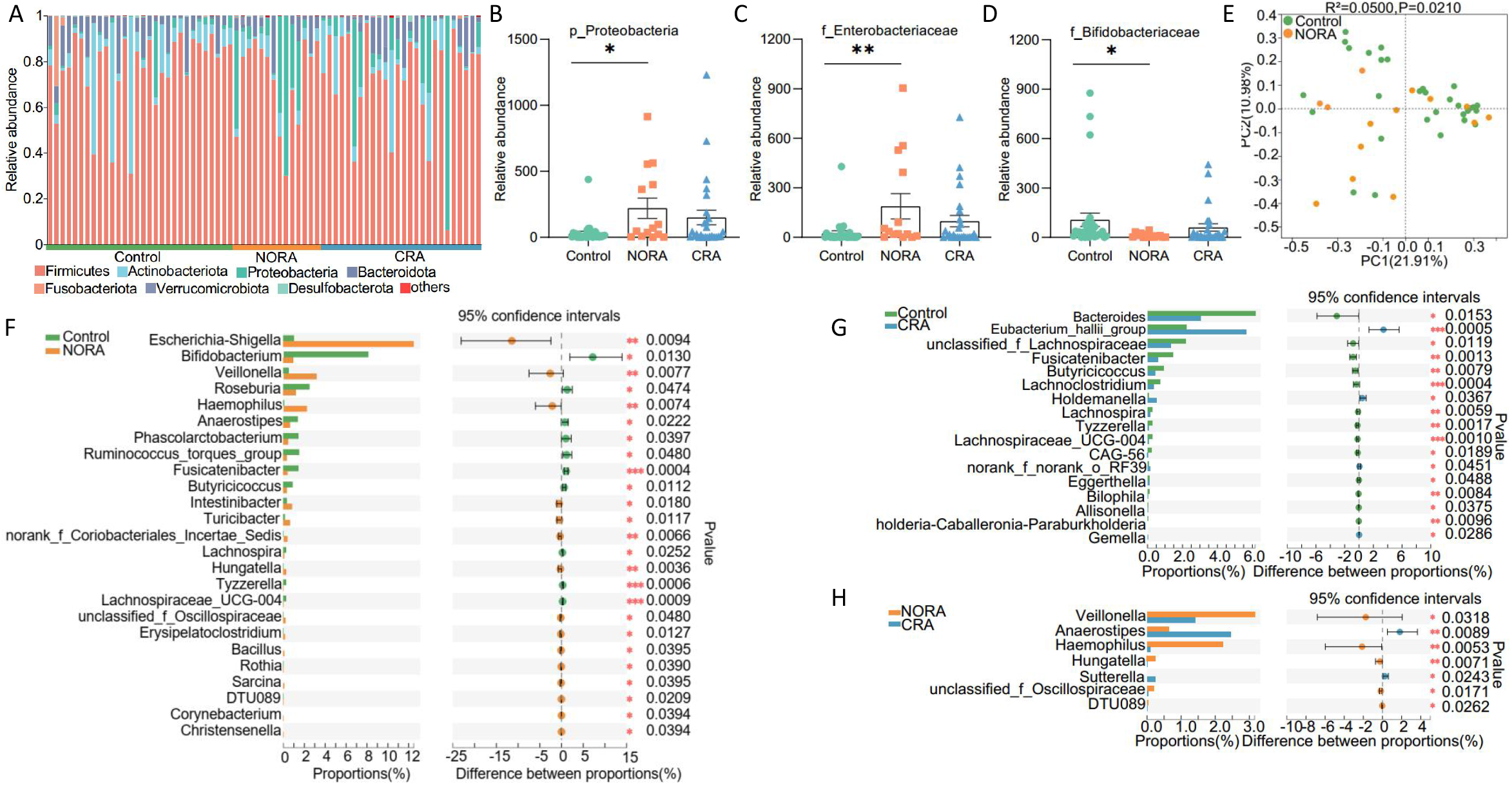
Alterations in the structural composition and diversity of the intestinal bacterial community. (A) Stacked bar graph showed the community composition at the phylum level among control, NORA and CRA. (B) Bar plots displayed the relative abundance expression of P_Proteobacteria in three groups, indicating a significant difference in abundance between new-onset RA and control. (C-E) The bar graphs showed the relative abundance of f_Enterobacteriaceae and f_Bifidobacteriaceae among the three groups, respectively. (E) PCoA revealed differential community structure between NORA and control in the beta diversity analysis. (F-H) Using the Wilcoxon rank-sum test, we identified 26 differential genera between NORA and control, 17 differential genera between CRA and control and 7 differential genera between NORA and CRA.

Linear discriminant analysis (LDA) demonstrated the importance of species from phylum to genus level among control, NORA, and CRA groups, we found that *f_Bifidobacteriaceae, g_Bifidobacterium, o_Bifidobacteriales* and *c_Actinobacteria* were dominant in control, while *p_Proteobacteria, g_Escherichia-Shigella* and *g_Veillonella* were predominant in NORA. Compared to CRA, *g_Bacteroides* and *f_Bacteroidaceae* were the predominant genera in control, and LDA analysis demonstrated the significance of *f_Pasteurellaceae* and *g_Anaerostipes* in both groups in NORA versus CRA (Supplementary Figure 3I-K).

These results indicated a significant variation in the abundance of gut bacteria from phylum to genus level between the three groups.

### Altered diversity of intestinal fungi among control, NORA, and CRA groups

To observe the alterations in intestinal fungal diversity and composition between control, NORA and CRA groups, we performed ITS sequencing on stool samples from subjects. Venn diagram showed a total of 64 species across three groups and 14 species between NORA and CRA patients (Supplementary Figure 4A). In alpha-diversity analysis, we found a measurable increase in community richness in ace, chao and sobs indexes for NORA compared to control and CRA. The diversity of shannon and pd indexes were obvious reduced in CRA compared with NORA (Figure 4A-E). We found that the abundance of *Aspergillaceae* was significantly higher in control than in RA patients as the dominant genus, while *Saccharomycetales_fam_Incertae_sedis* was the dominant genus in RA at family level with community composition analysis (Supplementary Figure 4B). The abundance of *Cladosporiaceae* was significantly reduced in RA patients compared to control, and the abundance of *Phaffomycetaceae, Debaryomycetaceae,* and *Didymellaceae* was increased in NORA (Supplementary Figure 4C-F). PCoA analysis demonstrated that control and CRA could be distinguished from each other, whereas NORA could not be distinctly separated from control and NORA from CRA (Figure 4F, Supplementary Figure 3G-H). Subsequently, using the Wilcoxon rank-sum test, we identified 17 differential genera between control and NORA, and 14 genera were differed in control and CRA groups, with an apparent decrease of *Cladosporium* in RA. *Candida* was significantly increased in CRA compared to control. In addition, we also found 8 differential genera between NORA and CRA (Figure 4G-I).

**Figure 4.**
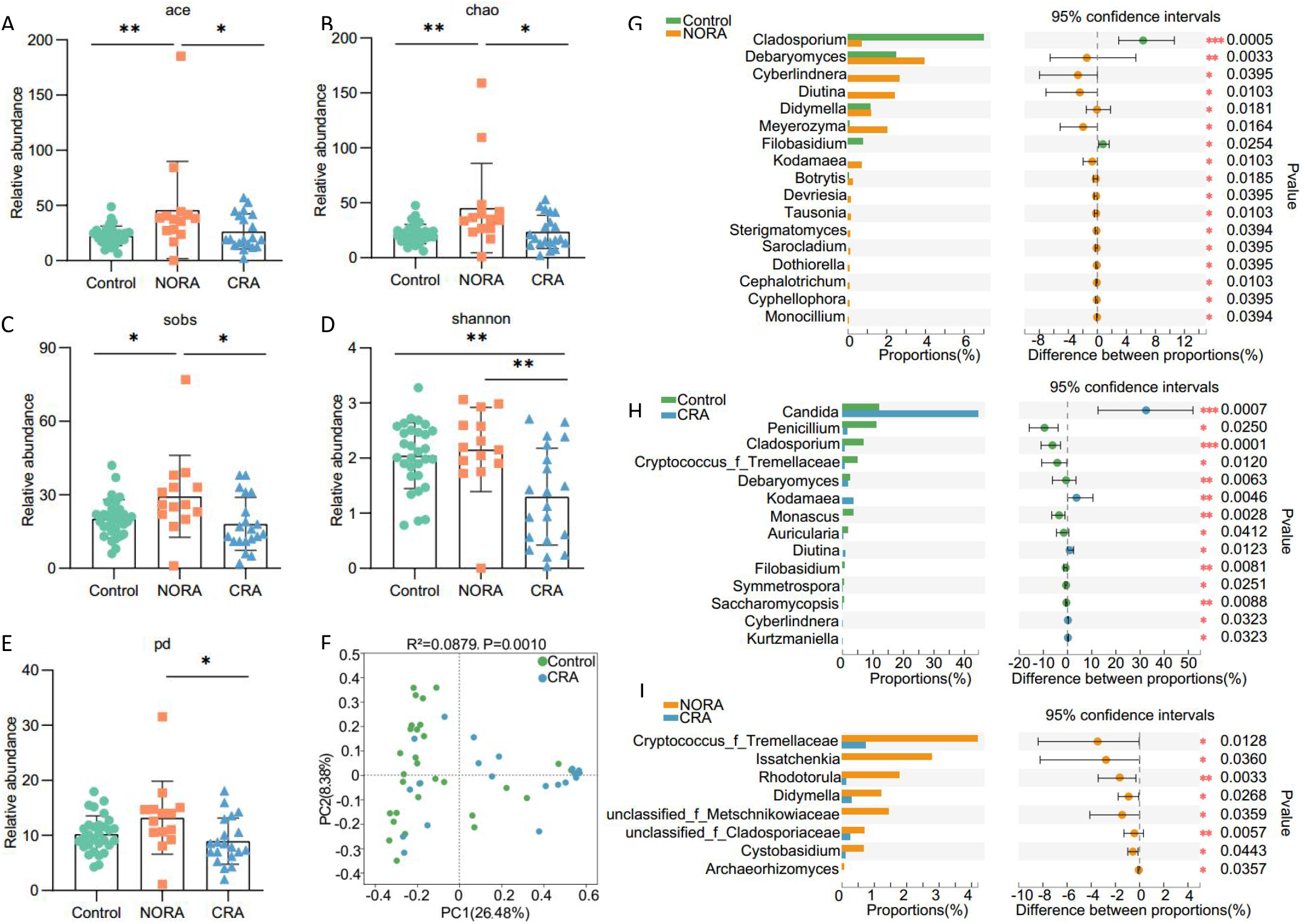
Changed intestinal fungal community composition and reduced diversity in CRA patients. (A-E) The bar graphs displayed the relative abundance of ace, chao, sob, shannon, and pd indexes between the three groups, respectively, with significantly lower community richness and diversity in CRA. (F) PCoA demonstrated significant differences in community structure between CRA and control. (G-I) Using the Wilcoxon rank-sum test, we identified 17 differential genera between NORA and control, 14 differential genera between CRA and control and 8 differential genera between NORA and CRA.

Taken together, these data suggested that dysbiosis of intestinal fungi was closely related to the progression of RA.

### Gene-expression profiles were significantly dysregulated between control, NORA and CRA patients

To further explore the characteristics of gene expression profiles in control, NORA and CRA, we performed RNA sequencing of PBMCs from subjects. Among NORA versus controls, we selected protein-coding genes for Gene Set Enrichment Analysis (GSEA), of which 9 gene sets were significantly enriched in control, including glycerophospholipid metabolism, glycosylphosphatidylinositol gpi anchor biosynthesis, calcium signaling pathway, taste transduction, neuroactive ligand receptor interaction, RNA polymerase, arachidonic acid metabolism, hedgehog signaling pathway, and basal cell carcinoma (Figure 5A, Supplementary Figure 5A-H, Supplementary Table 3). Furthermore, of the protein-coding genes we screened in CRA versus control, GSEA analysis showed 3 gene sets enriched in control and 16 gene sets enriched in CRA, like citrate cycle (TCA cycle) (Figure 5B, Supplementary Table 4). Subsequently, we identified 350 differential genes between NORA and control, a volcano plot showed 32 significantly up-regulated and 318 significantly down-regulated genes (|FC|>2, P adj<0.05). Between CRA and control, we found 690 differentially expressed genes and volcano displayed 249 genes significantly upregulated and 441 downregulated (|FC|>2, P adj<0.05). Also, we identified 40 differential genes between NORA and CRA, and the volcano plot demonstrated significant upregulation of 10 genes and downregulation of 30 genes (|FC|>4, P<0.05) (Figure 5C-E). Moreover, we performed KEGG enrichment analysis on 350 genes and identified 33 pathways that were significantly dysregulated, and bubble diagram showed 14 differential pathways, mainly in metabolic pathway and signaling pathways.

**Figure 5.**
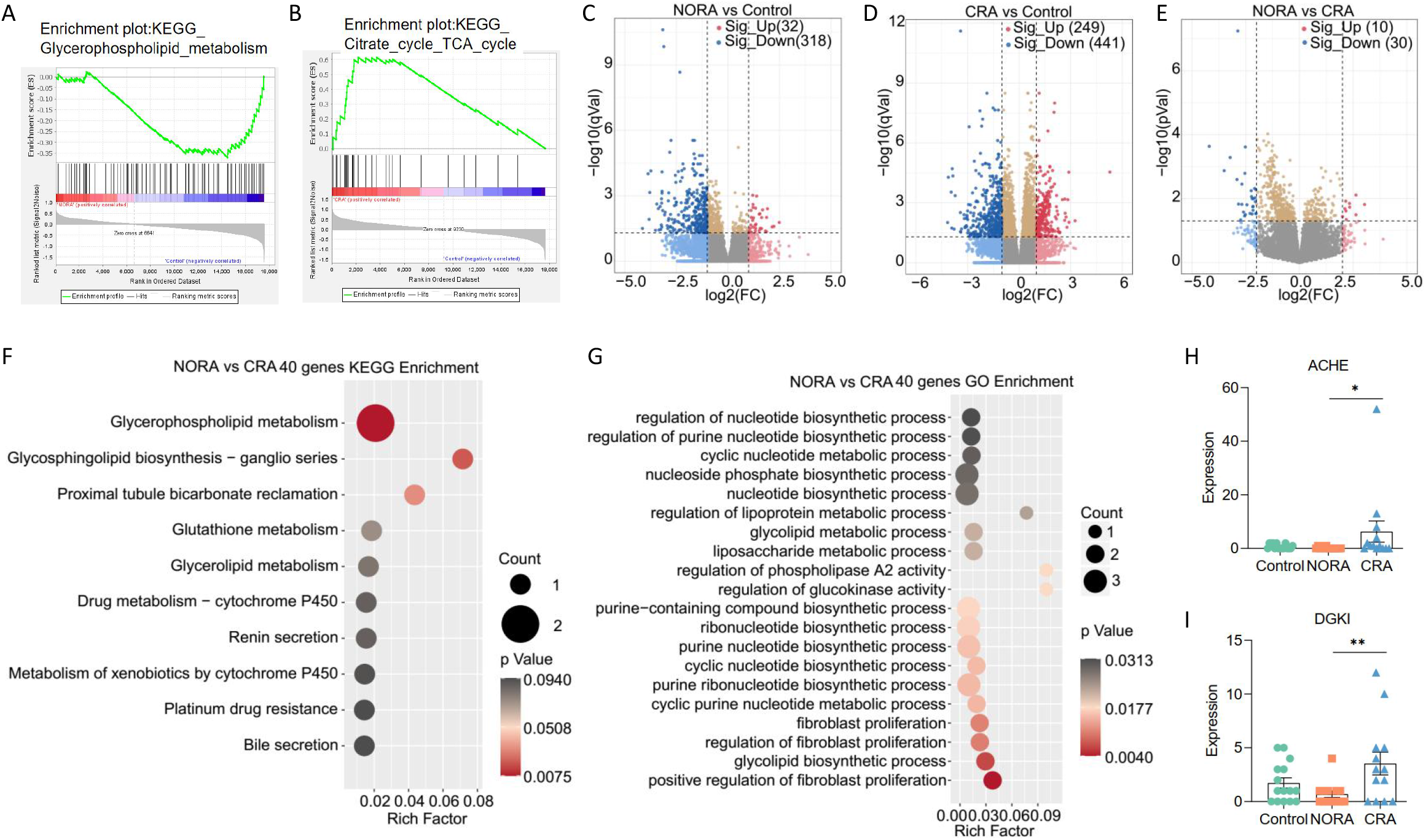
Differences in gene expression profile and enrichment analysis between NORA and CRA groups. (A) GSEA (Gene Set Enrichment Analysis) enrichment analysis demonstrated that glycerophospholipid metabolism was significantly enriched in the control compared to NORA group. (B) GSEA enrichment analysis demonstrated a significant enrichment of citrate cycle (TCA) cycle in CRA compared to control. (C) Volcano plot showed differentially expressed genes between NORA and control, with 32 upregulated and 318 downregulated genes. (D) Volcano plot showed 690 differentially expressed genes between CRA and control, with 249 up-regulated and 441 under-regulated genes. (E) Volcano plot showed 40 differentially expressed genes between NORA and CRA, with 10 were upregulated and 30 were downregulated. (F, G) Bubble plots illustrated the results of KEGG and GO enrichment analysis for 40 differentially expressed genes, respectively. (H, I) The bar graphs demonstrated the expression of ACHE and DGKI genes in the glycerophospholipid metabolism pathway.

We identified 364 pathways significantly disordered by Gene Ontology (GO) enrichment analysis, and the bubble demonstrated 21 of them, and we noticed that the differential genes were mainly enriched in type I interferon signaling pathway, cell differentiation, and G protein receptor signaling pathway (Supplementary Figure 5I-J, Supplementary Table 5). Based on 690 differentially expressed genes between CRA and control, we identified 128 pathways significantly enriched using KEGG enrichment analysis, with bubble displayed 15 pathways with significant dysregulation in arginine biosynthesis, histidine metabolism and glycerophospholipid metabolism. 784 pathways were discovered to be apparently altered in GO analysis, with bubble showed 26 of them (Supplementary Figure 5K-L, Supplementary Table 6). Also, we performed KEGG and GO enrichment analysis of 40 differentially expressed genes between NORA and CRA patients. Meanwhile, we identified 3 pathways significantly dysregulated, mainly enriched in glycerophospholipid metabolism with KEGG analysis. Using GO enrichment analysis, we identified 128 pathways of biological process significantly dysregulated, and bubble plot showed 20 of them, mainly enriched in biosynthetic and metabolic processes (Figure 5F-G, Supplementary Table 7). Furthermore, we found the expressions of ACHE and DGKI were significantly increased in CRA compared to NORA patients in glycerophospholipid metabolism pathway, which may be associated with dysregulated lipid metabolism (Figure 5H-I).

These outcomes indicated a significantly dysregulated gene expression profile in RA patients and a predominant enrichment in the glycerophospholipid metabolism pathway.

### Multi-omics combined analysis and ROC classification models establishment

To further explore the potential interrelationship between differential metabolites and genes, we focused on the interconnection between proteins in the glycerophospholipid metabolism and phenylalanine metabolism pathways enriched by 28 differential metabolites and 12 genes in the differential gene enrichment analysis. Protein-protein interaction network analysis demonstrated that acetylcholinesterase (ACHE), fibronectin 1 (FN1) and aquaporin 1 (AQP1) were core genes, and we found that ACHE was interlinked with lysophospholipase I (LYPLA1), AQP1 and FN1, and lecithin-cholesterol acyltransferase (LCAT) was interlinked with LYPLA1, glucokinase regulator (GCKR) and tyrosine aminotransferase (TAT). The ACHE gene was enriched in the glycerophospholipid metabolism pathway, and the downstream metabolites of LCAT and LYPLA1 proteins were the differential metabolites 1-Palmitoyl-sn-glycero-3-phosphocholine, 1-Stearoyl-2-hydroxy-sn-glycero-3- phosphocholine and glycerophosphocholine, suggesting that the LCAT and LYPLA1 proteins interact with the ACHE gene, resulting in altered metabolites in the glycerophospholipid metabolic pathway (Figure 6A). Correlation heatmap and network demonstrated strong correlation between differential flora, differential metabolites and clinical inflammatory indicators, and we found that CRP, DAS28, ESR and IL-6 were significantly negatively correlated with glycerophosphocholine, whereas positively correlated with the level of L-Phenylalanine. Trans-cinnamate showed a significantly positive correlation with CRP and negative correlation with *Sutterella.* DAS28, ESR and TJC28 showed a significant positively correlation with alpha-N-Phenylacetyl-L-glutamine. *Veillonella* was negatively correlated with 1-Palmitoyl-sn-glycero-3-phosphocholine, while positively correlated with *Hungatella* and *Haemophilus* genera (Figure 6B, Supplementary Figure 6A).

**Figure 6.**
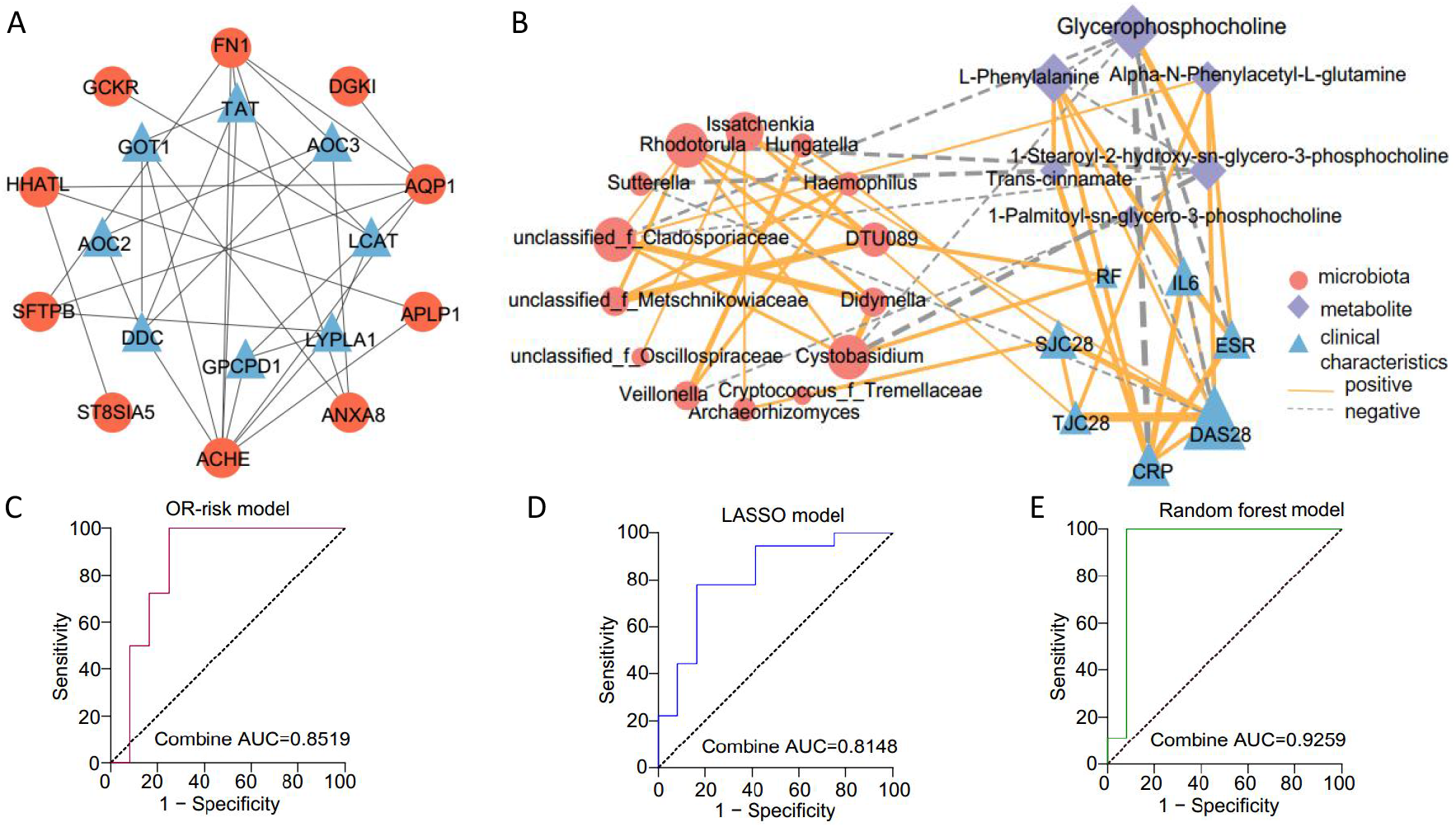
Multi-omics combined analysis and ROC classification models establishment. (A) Protein-protein interaction network analysis illustrated the interconnections between 12 differentially expressed genes and proteins on the glycerophospholipid metabolism and phenylalanine metabolism pathways between NORA and CRA. Red represented 12 of the 40 differentially expressed genes between NORA and CRA, and blue represented the proteins on the glycerophospholipid metabolism and phenylalanine metabolism pathways. (B) Correlation heatmap and network revealed the interrelationship between differential bacteria, differential fungi, differential metabolites and clinical inflammatory indicators, showing a strong correlation between flora, metabolites and inflammatory features. Red represented the differential flora, purple represented the differential metabolites, and blue represented the clinical inflammatory features. The size of the graph represented degree, the thick line of the line represented the correlation, the solid line represented the positive correlation, and the dashed line represented the negative correlation. (C) ROC analysis demonstrated a combined AUC of 0.8519 for the 4 features selected by calculating the OR value. (D) ROC demonstrated a combined AUC of 0.8148 for the 2 features selected by LASSO machine learning. (E) ROC demonstrated a combined AUC of 0.9259 for the 6 features selected by random forest. ROC: Receiver Operating Characteristic.

To identify crucial biomarkers to illustrate the differences between NORA and CRA, we selected 10 candidate markers with an area under the curve greater than 0.7 based on 7 differential bacteria, 8 differential fungi and 6 differential metabolites (Supplementary Figure 6A-C). Subsequently, we applied computational OR value, LASSO and random forest algorithms to identify essential features to build classification models to distinguish NORA from CRA patients, respectively. We filtered 4 key features (1-Stearoyl-2-hydroxy-sn-glycero-3-phosphocholine, glycerophosphocholine, *g_Rhodotorula, g_Cystobasidium)* by calculating OR value, and the AUC of the area under the curve was 0.8519. Meanwhile, the LASSO machine learning algorithm selected *g_Cystobasidium* and glycerophosphocholine with a combined AUC of 0.8148. The random forest algorithm screened 6 key features with an AUC of 0.9259 for the area under the curve (Figure 6C-E, Supplementary Figure 7D-F). These models demonstrated the good performance of three methods capable of distinguishing NORA from CRA patients.

The data suggested a potential relationship between intestinal flora, metabolites and clinical inflammatory indicators, which play an important role in the progression of RA.

## Discussions

Our findings explain the dysregulation in plasma metabolic profiles, gut flora and gene expression profiles between the control, NORA and CRA, revealing that the pathogenesis of RA involves multi-factor interaction and regulation. We found that the differential metabolites between NORA and CRA are significantly enriched in glycerophospholipid metabolism and phenylalanine metabolism pathways. Meanwhile, we also found that the dysbiosis in the abundance and diversity of intestinal flora in RA patients. Additionally, we found that the differential genes are enriched in lipid metabolism, amino acid metabolism, biosynthesis and metabolic processes. Lastly, correlation analysis suggested a strong association between flora, metabolites, genes and clinical features in RA development.

Currently, a growing body of evidence suggests that the occurrence and development of RA are intimately tied to intestinal flora dysbiosis, and experimental studies have shown that intestinal dysbiosis triggers arthritis in mouse models (18–21). Previous studies have confirmed that *Prevotella* is significantly increased in the intestine of early-diagnosed RA patients and activated the immune system and immune response, suggesting that *Prevotella* alterations appear to be a crucial factor in the pathogenesis of RA (12, 22, 23). The composition and diversity of intestinal flora were significantly reduced in RA patients, along with a decrease in the abundance of beneficial bacteria and an increase in the abundance of harmful bacteria, and it has been proved that probiotics can slow the progression of RA and lower levels of inflammatory factors (24). According to our findings, NORA patients had higher abundance of the bacteria *Escherichia-Shigella* and *Veillonella* and lower abundance of *Bifidobacterium,* which accelerated the progression of RA, in agreement with previous findings (12, 25, 26). However, some studies on Chinese RA patients have found a rise in *Lactobacillus* abundance during acute phase of RA, suggesting that the role of probiotics in the development of RA remains unclear (27, 28). We also observed that the abundance of *Veillonella* was reduced and differential in patients with CRA when compared with NORA, indicating that the inflammatory state may influence the change in the abundance of the flora. *Proteobacteria* were more prevalent and *Firmicutes* and *Bacteroidota* were less prevalent in RA patients, which was in line with previous research findings (11, 25). The gut bacterial community structure was similar between NORA and CRA patients, but β diversity revealed a substantial divergence between control and NORA patients, indicating that intestinal bacteria were distinct between RA and control. Intestinal fungi play an important role in the development of RA, and in our study, we discovered a significantly decreased abundance and diversity in patients with CRA compared to NORA, as well as significantly increased abundance of *Candida.* These findings are consistent with those of Harry Sokol et al. in patients with inflammatory bowel disease, which suggests that altered abundance of *Candida* correlates with inflammatory status (29). According to these findings, intestinal flora imbalance may have a role in the occurrence and progression of RA, and a rise in pathogenic bacteria and a decrease in probiotic bacteria are crucial factors in the disease’s development.

Previous studies, both in plasma metabolic profiles and in fecal metabolic analysis, have shown that RA patients have considerable alterations in metabolites and metabolic pathways (11, 25, 30). In our findings, disturbances in glycerophospholipid metabolism and phenylalanine metabolism pathways which are enriched by differential metabolites in RA patients, suggested that lipid metabolism and amino acid metabolism pathways apparently play a key role in the initiation and progression of RA. Consistent with our result, Die Yu. et al. also found dysregulation of glycerophospholipid metabolism and amino acid metabolic pathways in RA patients (11). More importantly, we also discovered a strong correlation between metabolites and clinical characteristics, with the metabolite glycerophosphocholine showing a significant negative correlation with CRP, while the metabolites L-Phenylalanine and trans-cinnamate showed a significant positive correlation with CRP. DAS28 and ESR showed significant positive correlations with metabolites L-Phenylalanine and alpha-N-Phenylacetyl-L-glutamine and negative correlations with glycerophosphocholine, which justified the close association of metabolites with CRP in previous studies, revealing that the activation of inflammatory factors during inflammatory states may influence the metabolic levels of metabolites (31). In addition, we noted that the clinical data was associated with intestinal flora, with *Cystobasidium* and *DTU089* genera showed demonstrating a significant positive correlation with RF, and DAS28 being positively correlated with *Issatchenkia* genus and CRP expression levels and negatively correlated with *Sutterella.* Correlation analysis revealed that plasma metabolites are significantly dysregulated in RA patients and correlate and interact with intestinal flora and clinical features.

Inflammatory factors such as IL-7, IL-6 and tumor necrosis factor play an essential role in the activation of the inflammatory response in RA (32–34). Our results showed that differentially expressed genes were enriched in the TGF-beta signaling pathway, the IL-17 signaling pathway, the MAPK signaling pathway, indicating that the inflammatory response pathway was disrupted in RA. A study reported that the MAPK signaling pathway was involved in cellular pathways in diseases such as RA, which was in accordance with our results (35). Previous studies have described the involvement of interferons in a number of autoimmune diseases, including RA and SLE. N Macías-Segura et al. showed that gene expression of type 1 interferon signaling was associated with autoantibody production in RA(36–38). Consistent with the results of our GO enrichment analysis, significant dysregulation of type 1 interferon, platelet degradation, cell differentiation and inflammatory response was demonstrated in RA patients. Importantly, we found that the results of KEGG enrichment analysis of differential genes coincident with the results of KEGG analysis of differential metabolites, implying gene and metabolite interactions.

One of the highlights of our study was the multi-omics combination analysis of the potential associations among control, NORA, and CRA patients. Nevertheless, we noted the limitations of our study. First, the small sample size of our study may limit the reliability and accuracy of our findings, which will need to be investigated in multicenter cohort and experimental studies to further validate our findings. Second, although our study elucidated the progression of RA from molecular layer to metabolic level, we need to gain insight into the intrinsic associations between multiple omics to reveal the essential role of glycerophospholipid metabolism in the development of RA.

In summary, comprehensive multi-omics analysis demonstrates that there is an inextricable link between control, NORA and CRA, that intestinal flora interacts with plasma metabolites, and that the differential core gene may influence changes in glycerophospholipid metabolism pathway. These findings suggest that glycerophospholipid metabolism is involved in the pathogenesis and development of RA from the molecular to the metabolite level and plays a significant role in RA pathogenesis, offering suggestions for early clinical diagnosis and therapeutic approaches.

## Acknowledgments

We are particularly grateful to the Department of Rheumatology and Immunology for their support and assistance with this project, to all authors for their dedication to this study, and to all patients who participated in this study.

## Supplementary methods

### Blood and stool sample collection

Morning fasting blood samples were collected from each participant and centrifuged at 3500 rpm for 10 minutes, followed by transferring the supernatant to a 1.5 ml EP tube and centrifuging at high speed for 12,000 rpm for 10 minutes. Finally, aspirated the supernatant into another EP tube and stored at −80°C in the refrigerator until use.

### DNA extraction, PCR amplification and Illumina sequencing of fecal samples

Total DNA extraction from microbial communities was performed according to the instructions of the E.Z.N.A.® soil DNA kit (Omega Bio-tek, Norcross, GA, U.S.), and the quality of DNA extraction was determined using 1% agarose gel electrophoresis, and DNA concentration and purity were determined using NanoDrop2000. 16S rRNA sequencing V3-V4 variable region PCR amplification primers were 338F (5’-ACTCCTACGGGAGGCAGCAG-3’) and 806R (5’-GGACTACHVGGGTWTCTAAT-3’) and ITS amplification primers were ITS1F (5’-CTTGGTCATTTAGAGGAAGTAA-3’) and ITS2R (5’-GCTGCGTTCATCGATGC-3’). PCR products were recovered using 2% agarose gels, purified using AxyPrep DNA Gel Extraction Kit (Axygen Biosciences, Union City, CA, USA), and sequenced using Illumina Miseq PE300 platform (Majorbio Bio-Pharm Technology Co. Ltd. (Shanghai, China).).

### Non-targeted LC-MS/MS analysis

100 μL of liquid sample was added to a 1.5 mL centrifuge tube, followed by 400 μL of extraction solution (acetonitrile: methanol = 1:1), mixed for 30 s and then extracted by low-temperature sonication for 30 min (5°C, 40 KHz), the sample was left at −20°C for 30 min, centrifuged at 4°C, 13000 g for 15 min, the supernatant was removed, blown dry under nitrogen, 120 μL of re-solution (acetonitrile: water = 1:1), low-temperature ultrasonic extraction for 5 min (5°C, 40 KHz), centrifugation at 13,000 g for 5 min at 4°C, and transferring the supernatant to the injection vial with internal cannula for analysis. In the on-line analysis, 1 QC sample was inserted in every 10 samples to observe the reproducibility throughout the analysis process. The analytical instrument was an ultra-performance liquid chromatography tandem time of flight mass spectrometry UPLC -TripleTOF system from AB SCIEX.

### Transcriptomic sequencing

PBMCs were obtained from 39 subjects prior to the administration of the drug, and RNA was extracted from the cells using standard extraction methods by company (Novogene, Beijing), and quality controlled and tested for RNA integrity using an Agilent 2100 bioanalyzer. The library was prepared using the NEBNext® Ultra™ RNA Library Prep Kit for Illumina®, and the library was sequenced using Illumina NovaSeq 6000 after passing the library test. Finally, reads were generated and genetically matched to the read data using the HISAT2 (v2.0.5) reference gene library. Data were pre-processed for bioinformatics analysis.

### Statistical analysis

The raw data of 16S and ITS were processed by quality control and noise reduction and uploaded to the Majorbio Cloud Platform (https://cloud.majorbio.com) of Majorbio Bio-Pharm Technology Co. Ltd. (Shanghai, China). The sequences after DADA2 noise reduction processing are commonly referred to as ASVs (amplicon sequence variants). The number of sequences per sample needs to be leveled for analysis of data results such as Alpha diversity, Beta diversity and species composition.

Raw data of non-target metabolism were log10 logarithmized using Progenesis QI (Waters Corporation, Milford, USA) software and uploaded on the Majorbio Cloud Platform (https://cloud.majorbio.com) for data analysis. Orthogonal partial least squares discriminant analysis (OPLS-DA) was performed using the R package ropls (Version1.6.2). Significantly different metabolites were selected based on the variable weight values obtained from OPLS-DA analysis (VIP>1) and Wilcox’ test p-values (p<0.05) to screen for significantly different metabolites, and were analyzed by the KEGG database (https://www.kegg.jp/kegg/pathway.html) and the Python package scipy.stats for metabolic pathway enrichment to obtain the pathways involved in differential metabolites.

We applied STRING for protein-protein interaction analysis, and the Spearman test was used for correlation analysis. In addition, we used SPSS 25 and Graphpad Prism 8 for data analysis.

**Supplementary Figure 1.**
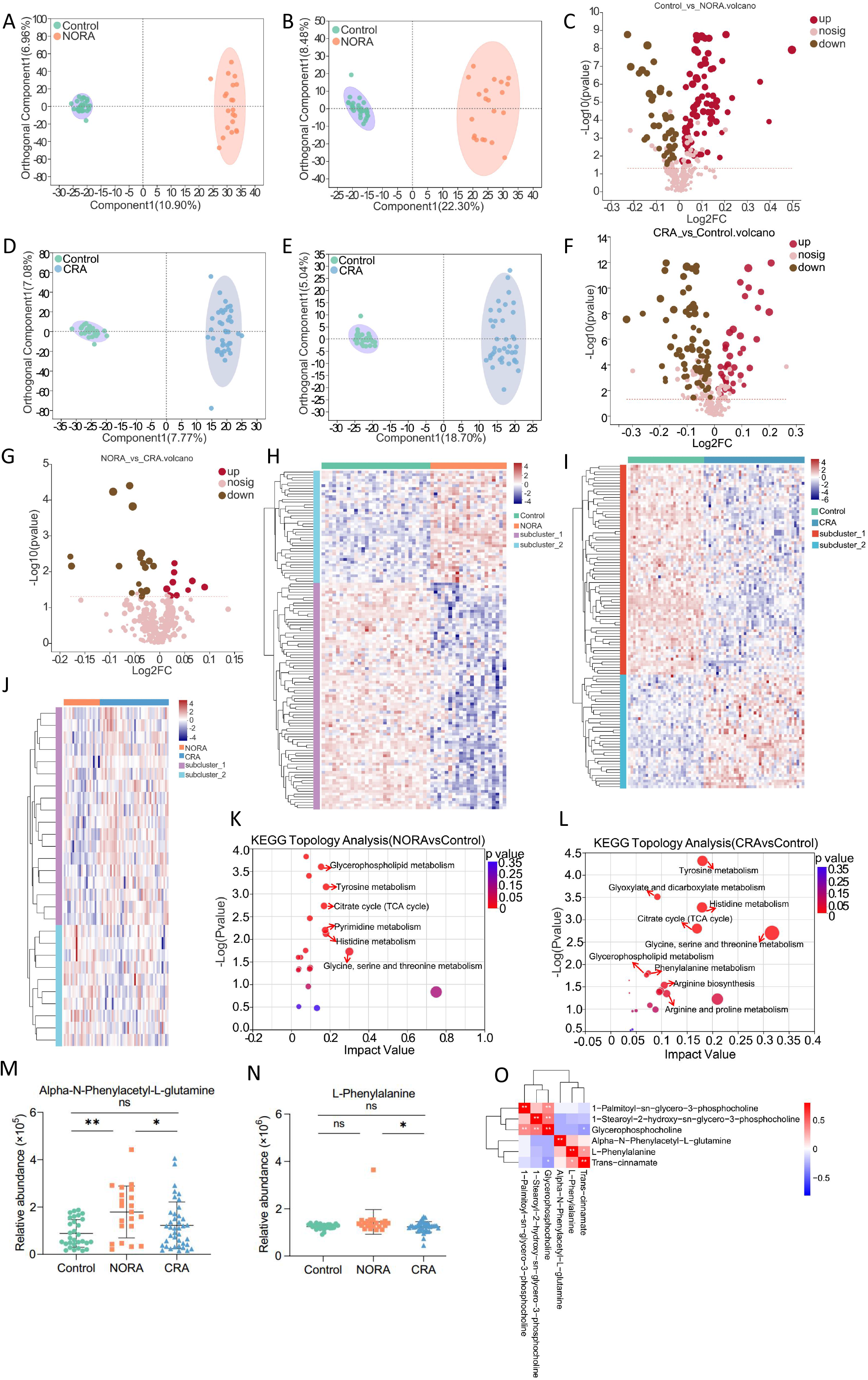
Analysis of plasma metabolic profiles between control, NORA and CRA patients.

**Supplementary Figure 2.**
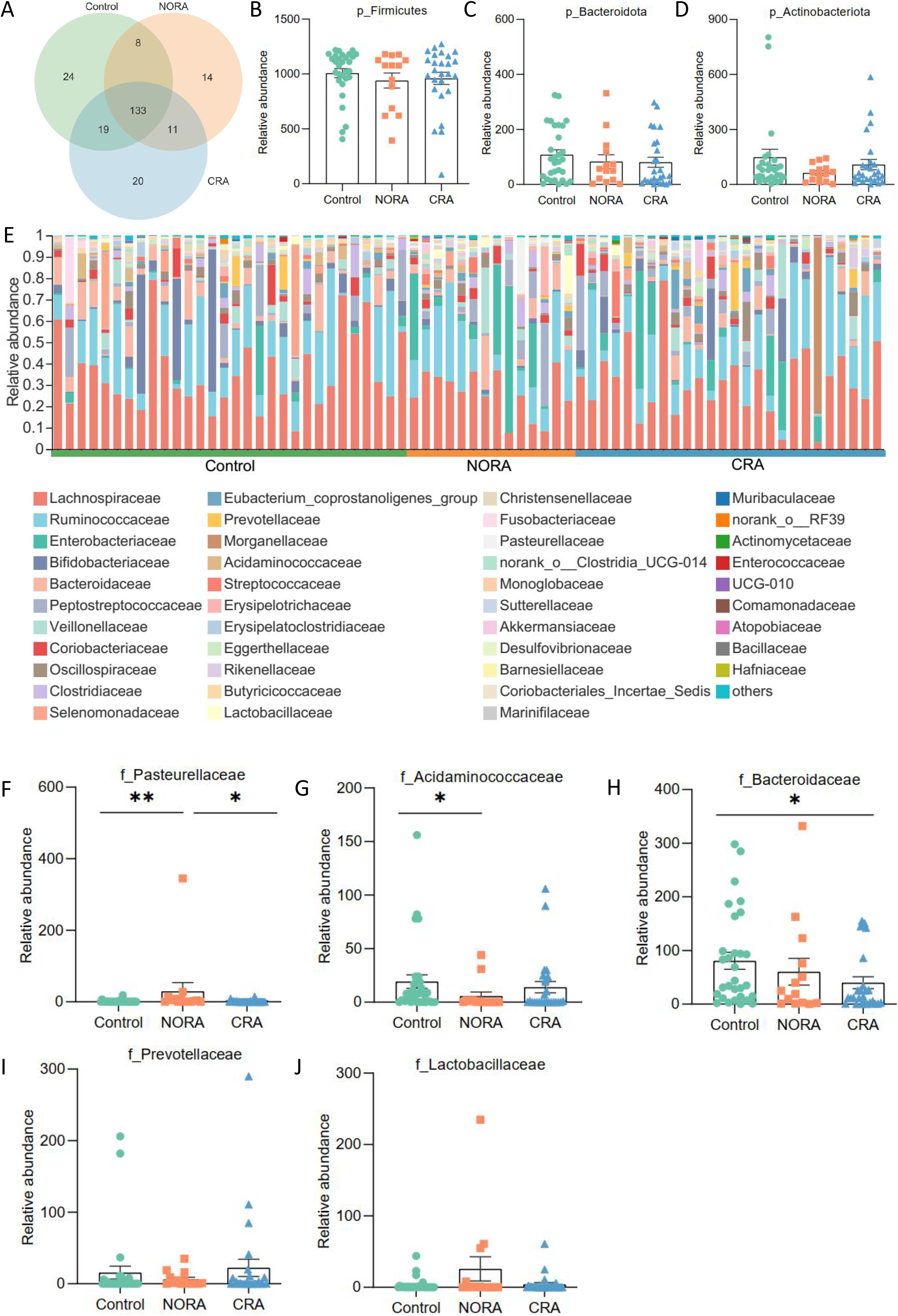
Gut bacterial community composition among control, NORA and CRA groups.

**Supplementary Figure 3.**
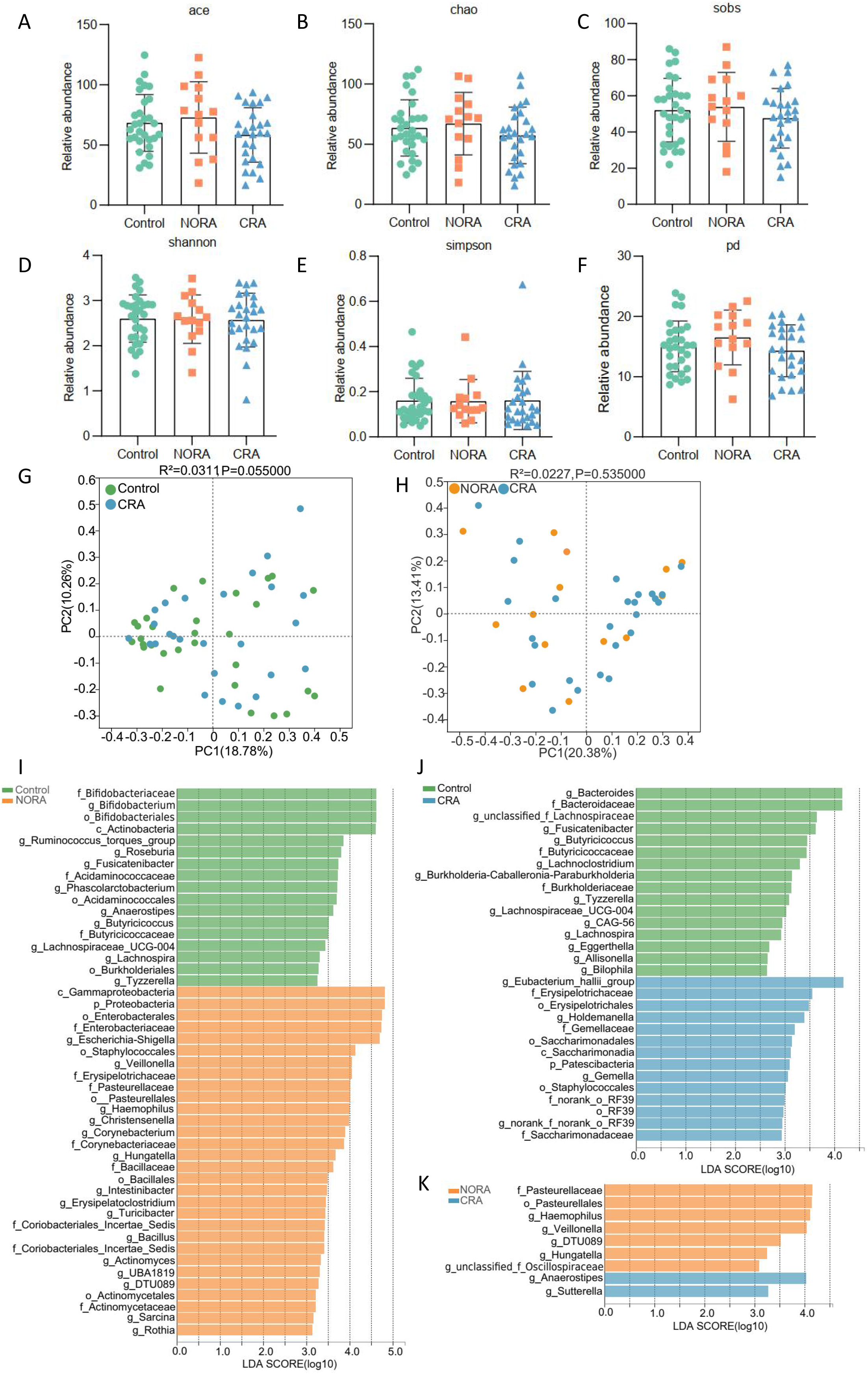
Gut bacterial community diversity among control, NORA and CRA groups.

**Supplementary Figure 4.**
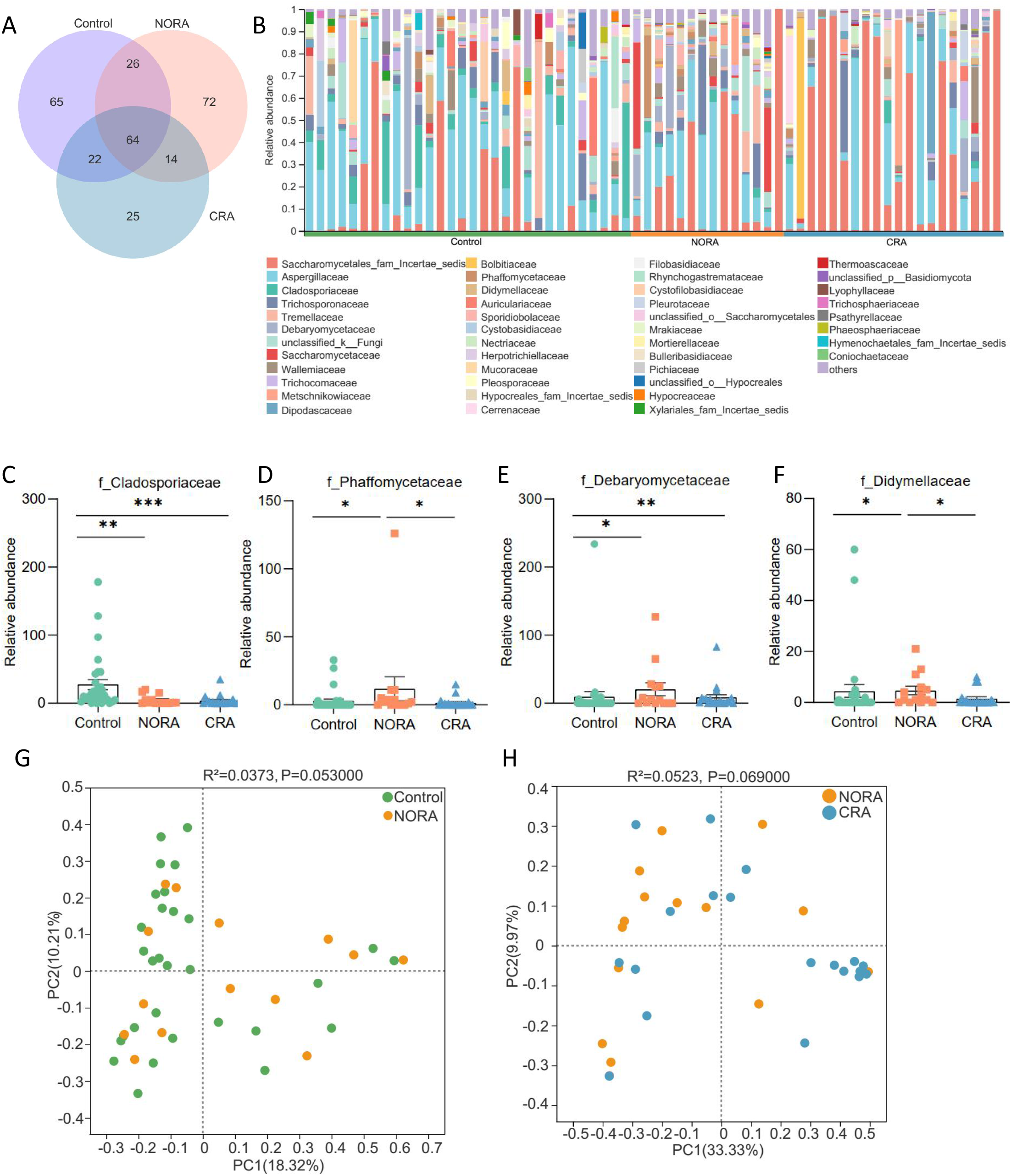
Intestinal fungal community composition and diversity between control, NORA and CRA groups.

**Supplementary Figure 5.**
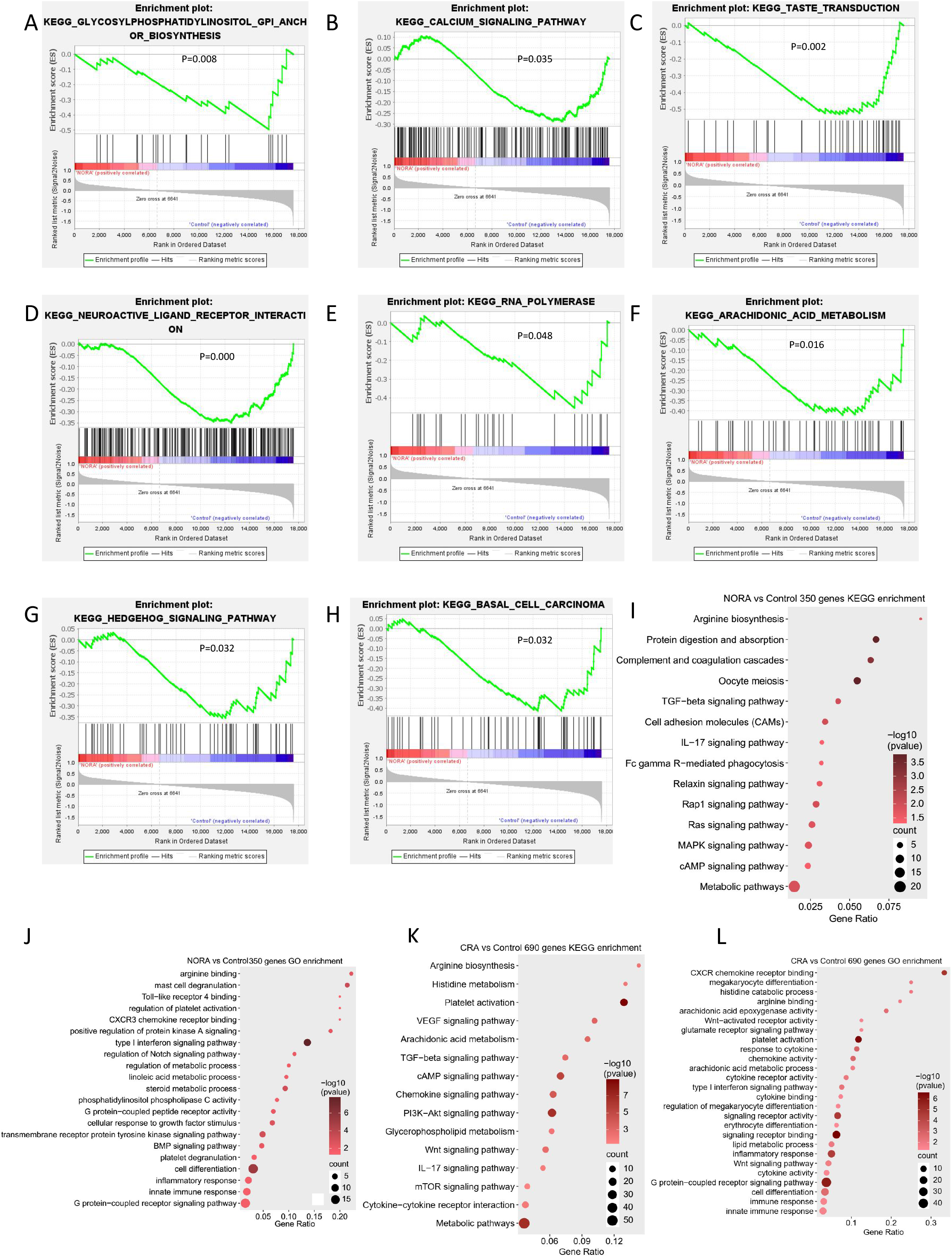
Gene enrichment analysis between control, NORA and CRA groups.

**Supplementary Figure 6.**
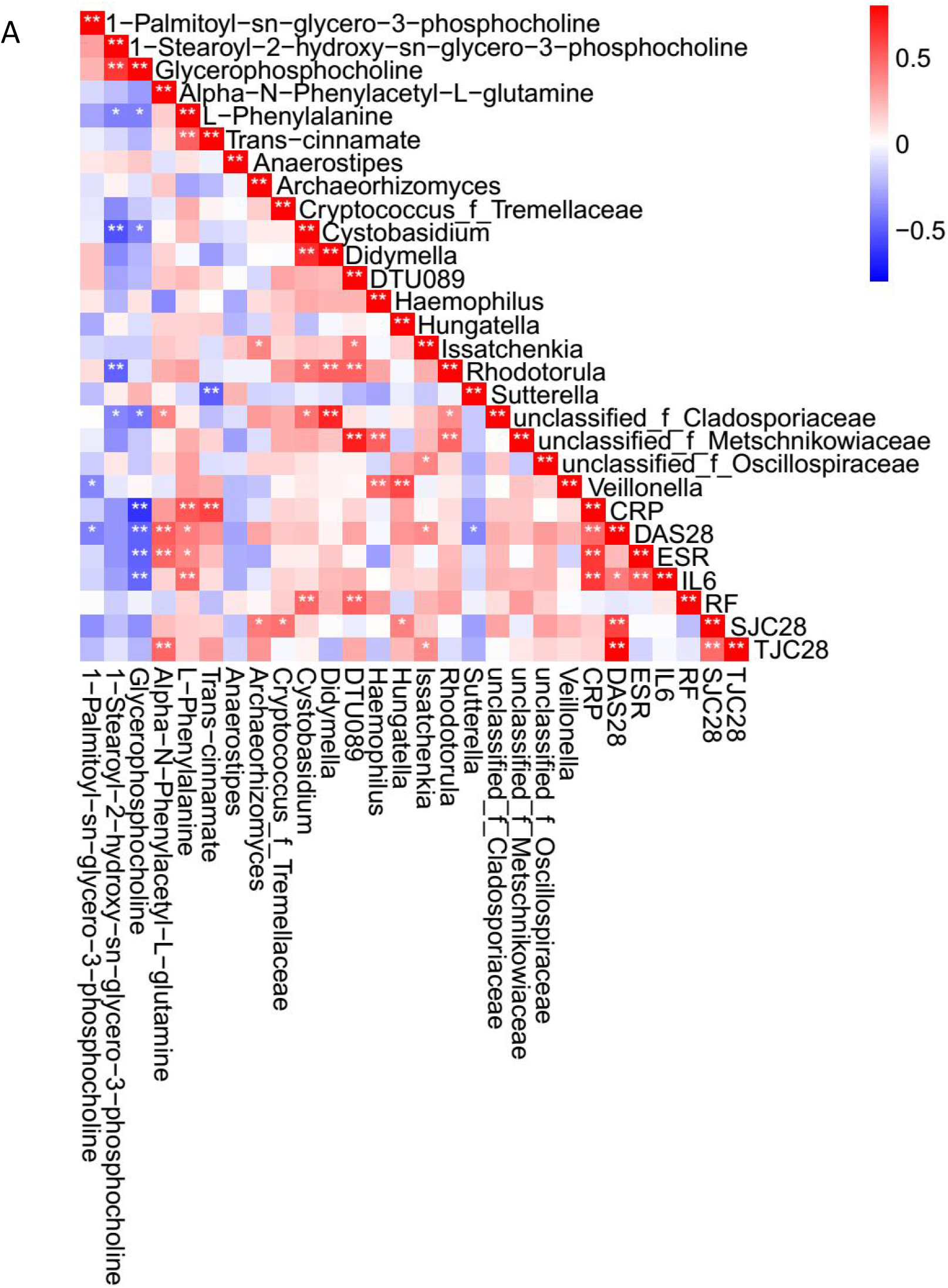
Correlation heat map showed the interactions and associations between differential flora, differential metabolites and clinical features.

**Supplementary Figure 7.**
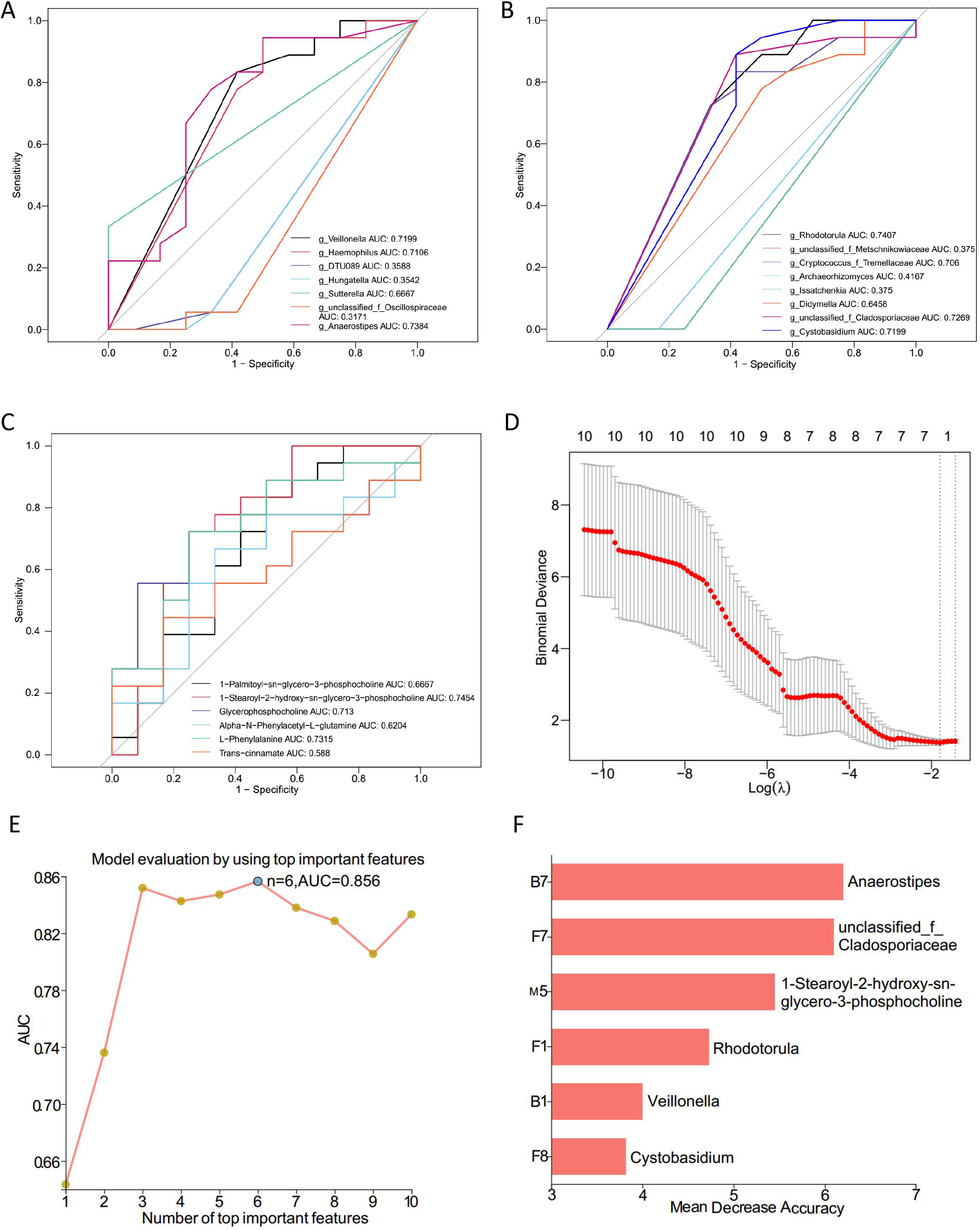
Signatures were selected based on LASSO machine algorithms and random forest.

